# A machine learning approach for the spatiotemporal forecasting of ecological phenomena using dates of species occurrence records

**DOI:** 10.1101/435289

**Authors:** César Capinha

**Affiliations:** CIBIO-InBIO, Centro de Investigação em Biodiversidade e Recursos Genéticos da Universidade do Porto, Campus Agrário de Vairão, R. Padre Armando Quintas, 4485-661 Vairão, Portugal; CEABN-InBIO, Centro de Ecologia Aplicada, Instituto Superior de Agronomia, Universidade de Lisboa, Tapada da Ajuda, 1349-017 Lisboa, Portugal.

**Keywords:** citizen science, GBIF, mushroom emergence, phenology, spatiotemporal forecasting, species occurrence records

## Abstract

1. Spatiotemporal forecasts of ecological phenomena are highly useful and significant in scientific and socio-economic applications. Nevertheless, developing the correlative models to make these forecasts is often stalled by the inadequate availability of the ecological time-series data. On the contrary, considerable amounts of temporally discrete biological records are being stored in public databases, and often include the sites and dates of the observation. While these data are reasonably suitable for the development of spatiotemporal forecast models, this possibility remains mostly untested.
2. In this paper, we test an approach to develop spatiotemporal forecasts based on the dates and locations found in species occurrence records. This approach is based on ‘time-series classification’, a field of machine learning, and involves the application of a machine-learning algorithm to classify between time-series representing the environmental conditions that precede the occurrence records and time-series representing other environmental conditions, such as those that generally occur in the sites of the records. We employed this framework to predict the timing of emergence of fruiting bodies of two mushroom species (*Boletus edulis* and *Macrolepiota procera*) in countries of Europe, from 2009 to 2015. We compared the predictions from this approach with those from a ‘null’ model, based on the calendar dates of the records.
3. Forecasts made from the environmental-based approach were consistently superior to those drawn from the date-based approach, averaging an area under the receiver operating characteristic curve (AUC) of 0.9 for *B. edulis* and 0.88 for *M. procera*, compared to an average AUC of 0.83 achieved by the null models for both species. Prediction errors were distributed across the study area and along the years, lending support to the spatiotemporal representativeness of the values of accuracy measured.
4. Our approach, based on species occurrence records, was able to provide useful forecasts of the timing of emergence of two mushroom species across Europe. Given the increased availability and information contained in this type of records, particularly those supplemented with photographs, the range of events that could be possible to forecast is vast.

## Introduction

Spatiotemporal predictions of ecological phenomena such as phenology, population dynamics and species interactions are fundamental to investigate the impact of climate change on future biodiversity (Urban et al., 2016) and to forewarn about risks for conservation (Franklin, 2010), human-health (Prank et al., 2013) and economy (Moriondo, Maselli & Bindi 2007). These predictions require the use of process-based or correlative models (Chuine & Regnier, 2017; Dietze, 2017). Process-based models involve experimental measurements under controlled settings but are too expensive to implement for many phenomena (Chuine & Regnier, 2017). Correlative models, however, relate non-manipulative ecological observations to putative environmental drivers, and ecologists generally find these more accessible. This approach underlies several recent examples of ecological forecasting (e.g. Scales et al., 2017), and has received significant methodological and conceptual advancements in recent years (e.g. Dietze, 2017).

The use of statistical models for spatiotemporal prediction in ecology remains strongly constrained by the availability of observational data. These data need to be spatially and temporally explicit, ideally consisting of long time-series recorded for various locations in the area under investigation (Jeanneret & Rutishauser, 2010). Data possessing such ‘ideal’ characteristics can be obtained via systematic field sampling or remote sensing (e.g. Jeanneret & Rutishauser, 2010; Moriondo et al., 2007; White, Thornton & Running 1997). However, the range of ecological events represented by these approaches remains limited. In light of this fact, it is perhaps relevant to consider the use of alternative data in model development.

One type of data that could suit this purpose is species’ occurrence records. These records provide the locations where organisms were observed and are now available in large numbers from formal (e.g. museum-records) and informal sources (e.g. geo-tagged photographs or video-based observations), for a wide range of taxa and across expansive geographical extents (Barve, 2014; García-Roselló et al., 2015). Unsurprisingly, these data play a key role in mapping and predicting the distribution of many species (e.g. ElQadi et al., 2017). A frequently more overlooked feature of these data is that it also often includes the dates of when the observations were made. Recent studies reveal that dates in the occurrence records can in fact describe the timing of the ecological events, such as pollinator species activity (Balfour, Ollerton, Castellanos & Ratnieks 2018; Bishop et al., 2013), mushroom fruiting phenology (Andrew et al., 2018) or plant flowering (Chapman, Bell, Helfer & Roy, 2015). Accordingly, correlative models fitting the temporal variation in species observation records across space may be useful for predicting the spatiotemporal dynamics of ecological phenomena.

In this work, we describe a framework to build temporal predictions of ecological phenomena across space using dates of species occurrence records. Our approach is framed within the context of ‘time-series classification’ (Geurts, 2001), a field of machine learning whose goal is to classify time-series into two or more classes. In simple terms, the approach aims to distinguish between time-series of environmental drivers that are related to the observation of the phenomenon and those that are not. We demonstrate the application of this approach using occurrence records for two mushroom species, *Boletus edulis* (‘the Cep’) and *Macrolepiota procera* (‘the Parasol’), and test if temporal variation in temperature and precipitation can predict the observation of their fruiting bodies across Europe. We used observations collected from conventional (e.g. museum records) and less conventional (e.g. geo-and time-tagged photographs) types of occurrence data, illustrating the potential of application to events captured by both types of sources. We observed that this approach consistently outperforms the predictive accuracy that other models have achieved using calendar dates and geographical coordinates, a standard ‘null’ model in spatiotemporal ecological prediction. We discuss further potential applications of the modelling approach and possible ways of improving it in the future.

## Materials and Methods

### Time-series classification for temporal prediction of ecological events

Time series classification uses machine-learning algorithms to classify time series into predefined category sets. Among several applications, the time-series classification has been used to detect ‘normal’ or ‘abnormal’ heart rhythms in ElectroCardioGrams (Kampouraki, Manis & Nikou, 2009) and to identify insect species from the frequencies of their wing-beats (Potamitis, Rigakis, & Fysarakis 2015). The classification can be done using the ‘raw’ time series or a set of predictors which summarise their properties (i.e., the ‘features’, in machine learning parlance) (Schäfer & Leser, 2017). The ‘raw’ series approaches calculate a point-by-point similarity with the time series of known classes. However, this approach is not very accurate when long or noisy time series are used (Fulcher & Jones, 2014; Schäfer, 2015) and thus they may be limited in their use in ecology (Hsieh, Anderson & Sugihara, 2007). Feature-based approaches, on the contrary, work by summarising the time-series into features. The objective of these features is to decrease the dimensionality of the raw data while retaining the information pertinent for classifying the data. The workflow of the feature-based approaches include: 1) collecting the time-series from the distinct classes; 2) transforming the time-series into features; 3) fitting a classification algorithm using the features as predictors; 4) evaluating the predictive accuracy of the model, and checking whether adequate accuracy levels are achieved and, 5) utilising the model to classify a new time-series.

This workflow bears similarity to many other prediction exercises in ecology. The only step that should be slightly unfamiliar to most modellers is the transformation of the time-series into features. Here two, non-mutually exclusive options are available – either to use a fully automated transformation such as discrete wavelet transform or discrete Fourier transform (e.g. Mörchen, 2003), or to extract the properties based on ‘expert knowledge’; Bagnall, Lines, Bostrom, Large & Keogh, 2017). One well-known example of a transformation based on expert knowledge involves the calculation of growing degree-days (i.e., the ‘feature’) from time-series of temperature, in order to achieve a more proximal representation of the effect of the accumulated heat on the plant and animal development (e.g. Neuheimer & Taggart, 2007).

### Spatial time series classification

By definition, time-series classification is concerned with temporal data. However, predicting the timing of the ecological phenomena is often a necessity for multiple locations. Importantly, different locations can imply different ecological responses to the same drivers, due to the influence exerted by the temporally invariant factors (like soil and land-use types) or to local adaptations of species and communities (Almeida‐Neto & Lewinsohn, 2004; Chuine & Regnier, 2017). Suitably, the time-series classification can be extended across space merely by including features representing the spatial dimension into the model. One example of the way this can be achieved is by using the *x* and *y* coordinates of the observations as features. Another approach, perhaps more inclusive, is to use eigenvectors from spatial connectivity matrices (see Griffith & Peres-Neto, 2006 for a description of the method).

### Case study

In this work we demonstrate the use of spatial time-series classification to predict the occurrence of the fruiting bodies of two mushroom species, *Boletus edulis* and *Macrolepiota procera,* in the countries in Europe. These species are collected and consumed by humans and are also a dietary component for wild fauna (Mazurkiewicz & Podlasińska, 2014). Spatially and temporally-explicit predictions of the occurrence of fruiting bodies of these mushrooms are thus arguably useful in managing their seasonal supply.

### Occurrence data

Records of the occurrence of the fruiting bodies of these species were collected from online sources. For practical purposes, these sources are distinguishable into those based on photographic records and those lacking such data. Photography-based records were collected from Flickr (https://www.flickr.com/), Mushroom Observer (https://mushroomobserver.org/), Observation.org (https://observation.org/) and Project Noah (https://www.projectnoah.org). Only photographs revealing the typical morphological traits of the species were considered. These traits for *M. procera* included a large white-to cream-coloured cap with brown scales and snakeskin markings on the stem, while for *B. edulis* it included a stem with an enlarged base and netted pattern. The photographic records also needed to include the day, month and year of its observation and location. Location could imply geographical coordinates or the name of a locality or region. The names provided were identified using Google Earth Pro (https://www.google.com/earth/index.html) and converted into geographical coordinates. Records with location names that were unidentifiable or having less than 10-km spatial accuracy were not considered.

We also collected occurrence records from the Global Biodiversity Information Facility (GBIF) (downloads https://doi.org/10.15468/dl.t39jw0 and https://doi.org/10.15468/dl.8juvkr for *Boletus edulis* and *Macrolepita procera*, respectively). From these we retained only those having complete date and geographical coordinates. As these records were not supplemented with photographs, some could refer to the non-fruiting forms of the species, such as its mycelium. To assess the possibility of this happening, we investigated the months in which the observations were made. We found that virtually all the observations had been done in months typical of the fruiting season of the species (Supporting Information Figure S1) – i.e., late summer and autumn, and less frequently in the spring. This result suggests that, if observations of the non-fruiting bodies are also included in the data, these should be limited in number and thus unlikely to affect the models.

Due to the meagre availability of records in many regions prior to 2009 and to allow time for observations to be added to the data sources, our analysis included a seven-year period, from 2009 to 2015. The data showed strong spatial bias, with most of the records coming from Scandinavia, Germany or Great Britain (Supporting Information Figure S2). For the other regions, mostly in the south fewer records were available, and were particularly ones that chiefly originated from photography-based data-sources (Supporting Information Figure S2). To minimise the spatial bias present in the data, which could overshadow the conditions sampled for regions with lesser number of records (Zadrozny, 2004), we down-sampled the number of records in some regions. The down-sampling was done by initially covering the study area with a grid of 200 × 200 km squares and counting the number of records in each square. The squares identified as upper outliers (i.e., number of records > Q3 + 1.5 × IQR), were down-sampled by randomly selecting a number of observations equal to the Q3 + 1.5 × IQR, where Q1 is the lower 25% quantile, Q3 is the upper 25% quantile and IQR = Q3 − Q1. We finally obtained a total of 2,441 observation records for *B. edulis* and 1,169 for *M. procera*. While these records were relatively well distributed over the years (Supporting Information Figure S3), both the species had fewer records in 2009 and a greater number in 2014.

### Temporal drivers and feature extraction

The potential occurrence of fruiting bodies of each species was classified using time-series of temperature and precipitation. While other factors, such as soil and habitat type could also affect mushroom fruiting, the temporal variability variabilities in temperature and water availability are strong predictors of mushroom fruiting (Diez, James, McMunn & Ibáñez 2013) and should allow capturing much of their temporal regularities. The meteorological data from Agri4Cast (http://agri4cast.jrc.ec.europa.eu/) were collected, which are available for European countries on a daily basis at 25×25 km resolution. The mean air temperature (°C) grids, as well as those of the accumulated precipitation (mm) were collected for every single day from 2009 to 2015. The two variables were temporally ordered and stacked into raster time-series.

We calculated a comprehensive set of features (*n* = 40; Supporting Information Table S1) to characterise each of the occurrence records in terms of the preceding values of temperature and precipitation. These features included, among others, the means of temperature and sums of precipitation for a diverse range of time windows. The length and limits of the time windows were adjusted to capture the detailed short-term meteorological variations (e.g. preceding weeks) and the more general variations in the mid-to long-term (e.g. preceding trimesters, semesters and the entire year). We observed that, as mentioned above, other approaches to transform the time series into features could have been employed. For a comprehensive review of automated transformations refer to Fulcher and Jones (2014).

To account for possible location‐dependent responses to meteorological variation, we also included the geographical coordinates (latitude and longitude) as features in the models. As mentioned earlier, more sophisticated means could have been used (e.g. Griffith & Peres-Neto, 2006), but this would have added an unnecessary layer of complexity to our illustrative aim.

Time series classification is generally done using data for two or more classes– but see Ma and Perkins (2003) for a one-class implementation. For our case study, the classes intuitively correspond to the ‘presence’ or ‘absence’ of fruiting bodies. However, no data is available on the ‘temporal absence’ of fruiting bodies of the species. Therefore, we contrasted the conditions represented by the occurrences with the range of conditions available to the species. More specifically, the models attempted to identify the subset of meteorological combinations related to the occurrence of mushroom fruiting from the entire set of combinations under which the species occurs. Sampling of the available conditions was done by randomly selecting a number of dates (in the 2009 to 2015 time range) for each occurrence record. These random records, tentatively termed ‘temporal pseudo-absences’ were then used to extract an equivalent set of features, referring to the location of the originating occurrence record. To accomplish this, we used 15 temporal pseudo-absences for each occurrence record. In preliminary models, this number provided a good balance between the comprehensiveness of the sampling and computation time required to run the models. For new data, this number is worth investigating.

Processing of the raster time-series and feature extraction was made in R, mainly utilising the functions from the raster package (v. 2.6-7).

### Implementations of the spatial time series classification model

We used boosted regression trees (BRT; Elith, Leathwick & Hastie, 2008) to classify between the occurrence of fruiting bodies and temporal pseudo-absences. Boosted regression trees are ensembles of individual regression trees, in which the trees are added in sequence – each fitting the residuals of the earlier ones. This modelling technique also includes a stochasticity component, which aims at minimising the effect of spurious patterns, and improving the generality of the model fittings.

The BRTs were implemented using the routine ‘gbm.step’ of ‘dismo’ (v. 1.1-4) package for R. Three parameters are relevant for fine-tuning BRT models: learning rate, tree complexity and number of trees. The learning rate (*lc*) refers to the contribution (weight) of each tree in the ensemble, tree complexity (*tc*) controls the interaction order on the response being modelled and the number of trees (*nt*) determines the total number of trees to be included in the ensemble. Besides, it is also necessary to define the stochasticity component (or bag-fraction), which refers to the proportion of data that is made available to grow the trees at each step.

Following the recommendations of Elith et al., (2008) here we used a fixed bag-fraction of 0.5, meaning that 50% of the data were randomly drawn at each step. The optimal settings for the other three parameters were determined iteratively by measuring model performance for all the combinations of *tc* values of 1, 3 and 5, and *lr* values of 0.01 and 0.005. For each combination of values of *tc* and *lr* tested, the optimal number of trees (*nt*) was automatically determined by ‘gbm.step’. Model performance was evaluated using a 5-fold cross-validation procedure and the measure used was the area under a receiver operating characteristic curve (AUC) (Bradley, 1997). The use of AUC is of specific relevance because this metric is insensitive to differences of prevalence (i.e. the ratio between the classes), and our data is strongly imbalanced towards pseudo-absences (15 for each occurrence).

### Comparison to a null model

Useful predictions are those that can recommend favourable changes from the usual patterns of activity (Lowe et al., 2015), which in the case of the present study, are based on the ‘normal’ fruiting season of each mushroom species. For instance, mushroom pickers often used the harvest dates of the previous years as an indicator of the potential dates for future harvests (e.g. Lincoff, 2015). In this context, to assess the practical worth of the predictions from our framework, we compared its predictive accuracy to the one provided by a model fitting the dates of the records.

This ‘null’, date-based model uses four features to describe the occurrence and pseudo-absence records: the sine and cosine transforms of the dates plus the latitude and longitude of the records. Using two-dimension transforms enabled the appropriate expression of the circular nature of the dates, which cannot be represented using only a single dimension, such as Julian days. The addition of the geographical position is also essential to account for the regional differences in the fruiting seasons.

The predictive performances of the ‘full’ and ‘null’ models were compared using a *k*-fold cross-validation, where *k* corresponded to each of the years in the data (i.e. 2009 to 2015). This procedure corresponded to 1) the utilisation of data for all the years, except for one, to identify the combination of the model parameters providing the higher AUC (see the previous section), 2) employment of the model with optimal parameters to make predictions for the ‘out-of-sample’ year and 3) measurement of the agreement between the values predicted and those observed. Testing was done for each of the years and, to account for the stochastic nature of the BRT which might produce slight differences in the predictive accuracy of the models using the same data, the model training-testing cycle was repeated five times for each year. The agreement level achieved between the predictions and left-out observations was measured using the AUC.

To ensure that the measurements of accuracy represented the entire study area and time periods, the distribution of deviations between predicted and observed values were mapped. The deviations corresponded to the difference between the averages of the predictions of the five replicate models and the values of the observations of the test year.

We also evaluated the temporal ‘behaviour’ of the predictions from the ‘full’ and ‘null’ models by visualising the way the predicted probabilities of the occurrence changed over time. This was done by making the predictions once in every five days, from 2009 to 2015 for three locations in the area under study (Supporting Information Figure S4). These predictions were made using the BRT models trained with the data from all the years and using the combination of the parameters most often identified during cross-validation as optimal (Supporting Information Figure, Table S2).

## Results

Models based on the temperature and precipitation time-series (i.e., ‘full’ models) consistently outperformed the date-based (‘null’) models in predicting the occurrences and pseudo-absences of the fruiting bodies for the two mushroom species (Table 1). Considerable improvement in accuracy is evident in some years, with the AUC values showing 10% (or higher) improvement. This improvement occurs even when the null models generally provide what may be regarded as a good predictive accuracy (i.e., AUC > 0.8), suggesting their ability to precisely capture the ‘average’ season of the mushroom emergence. Both large and small deviations between predicted and observed values are observed across the study area and for all the years of study (Supporting Information Figures S5-S8), supporting the spatial and temporal representativeness of the AUC values obtained.

**Table 1.**
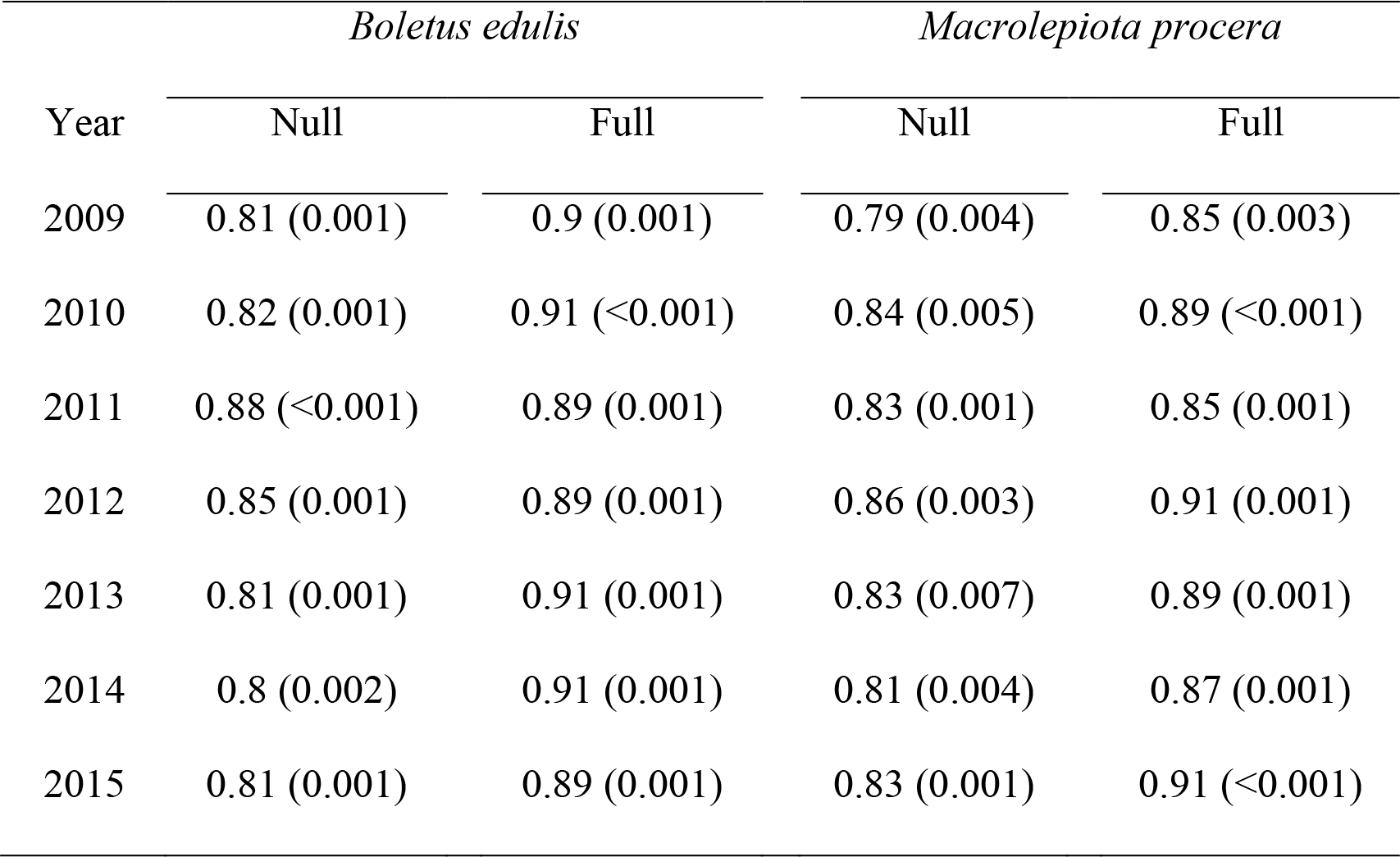
Accuracy of the boosted regression tree models (BRT) in predicting the occurrence of fruiting bodies of the mushrooms *Boletus edulis* and *Macrolepiota procera*. Two types of models are compared. In the ‘full’ models the occurrence and temporal pseudo-absence records are characterised in terms of the preceding environmental variations, while in the ‘null’ models the records are characterised using the calendar dates. Accuracy is measured using the area under the receiver operating characteristic curve (AUC) and refers to the ability of the models in predicting the observations for a year that is ‘left out’ of the model training. The testing is done for all the years, one year at a time.

Plots of the predictions made every five days compare the ‘average’ season of the mushroom emergence captured by the null models (Figure 1, Supporting Information Figure S9, grey area), with the environmental-driven responses of the full models (Figure 1, Supporting Information Figures S9, black line). These plots show substantial agreement between the two types of predictions, although for some years important differences can be observed. These differences include distinctly higher or lower ‘in-season’ probabilities of occurrence. For instance, in 2010, for a site in England (Supporting Information Figure S4), both species showed distinctly higher probabilities of occurrence from the environmental-based models than from the null models, while the inverse was true for 2011 (Figure 1). Seasonal lengths too showed differences. For instance, the environmental-based models predicted a shorter season for both species in 2012 and a longer season for *M. procera* in 2015 than did the ‘average’ calendar-based season (Figure 1).

**Figure 1.**
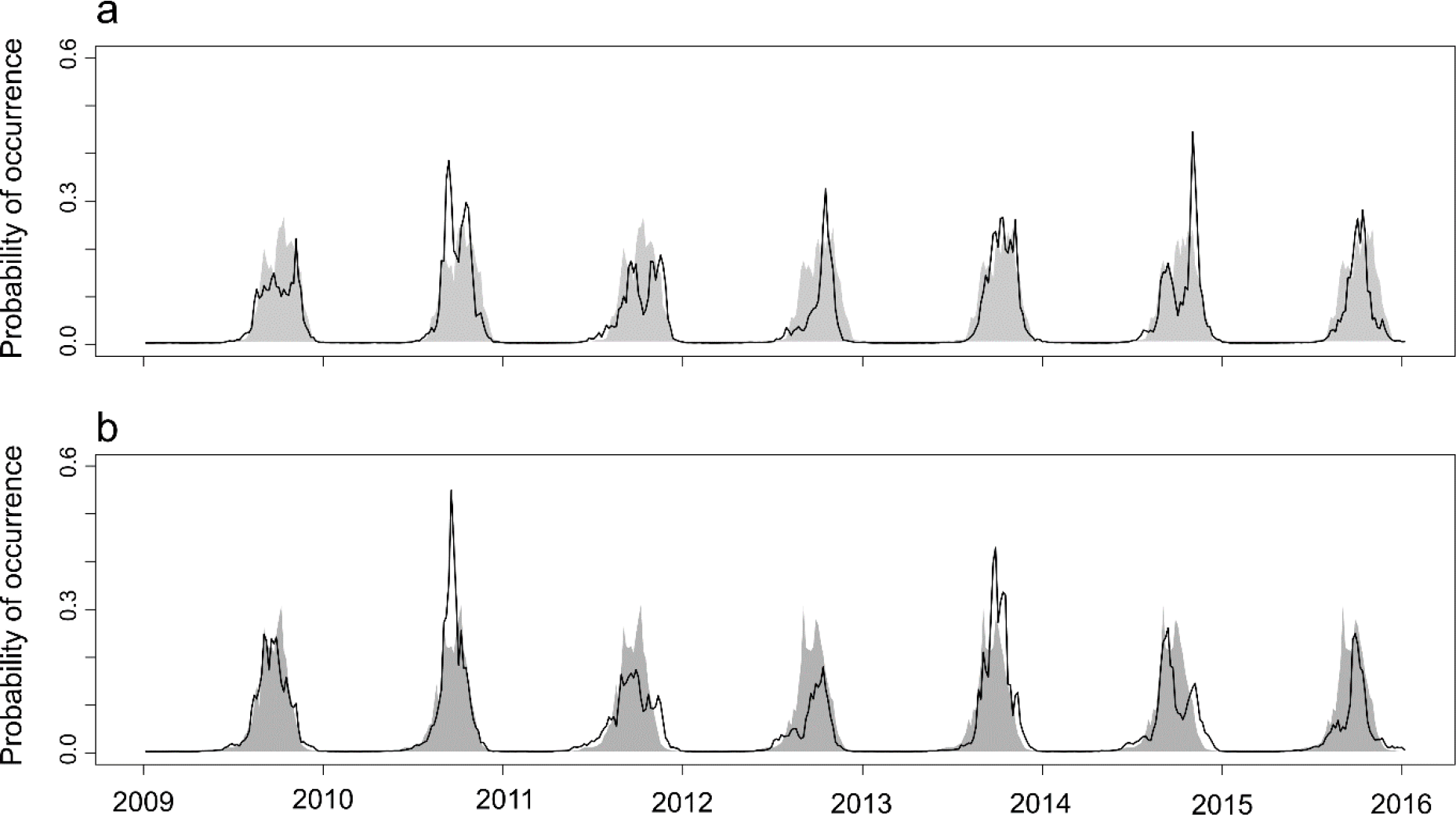
Predictions of the probability of occurrence of fruiting bodies for *Boletus edulis* (a) and *Macrolepiota procera* from 2009 to 2015, for a site near Cambridge, England.

Predictions are made at 5 days-intervals from 2009 to 2015 and compare a model trained with features that describe meteorological variation (black line; ‘full model’) and a model trained with features that describe the dates of the records (grey area; ‘null model’). A slight modification in the null model-predicted response after 2012 reflects a one-day date change caused by the leap year.

## Discussion

In this paper, we have described a machine learning approach to model the spatial and temporal information found in species occurrence records. We used the approach to model the timing of emergence of two mushroom species across Europe and found support for its use in forecasting emergence patterns in the future. We expect that this approach may be applied to forecast other phenomena represented by occurrence records.

Occurrence records that are supported by photographs or videos are specifically relevant to the application of the approach that we have demonstrated. The ecological phenomena that these records are able to capture are vast (e.g. leaf greenness, fruit ripening, adult insect activity, colour pelage in mammals etc.) and their availability in the public repositories continues to rapidly and extensively increase (e.g. Loarie, 2017). For instance, at the time of writing the present work, a search on Flickr using the expression ‘bee OR pollination’ retrieves more than one-and-a-half million records. This number will further escalate if additional data sources are considered, such as iNaturalist, Facebook and other social media sites and citizen-science projects. It is thus plausible to expect that the information drawn from these sources may prove adequate for the identification of relevant relationships between the spatiotemporal dynamics of many ecological factors and their drivers.

Our approach, as presented here, includes a basic set of conceptual and methodological guidelines, which could be expanded and improved upon in the future. A few methodological changes, in particular, may allow improving the predictive accuracy. One of these concerns the transformation of the environmental time-series into features. The transformation we employed was user-defined, aiming for simplification; however, there is strong support for the use of automated methods (e.g. Bagnall, Davis, Hills & Lines 2012), particularly for those that iteratively adapt the transformations to losses or gains in predictive performance (Flaxman, Chirico, Pereira & Loeffler 2018). Another possibility involves the mitigation of spatial and temporal bias in the data. These biases are a highly recognised pervasive characteristic of most data sets of biological records (Isaac & Pocock, 2015; Tiago, Ceia-Hasse, Marques, Capinha & Pereira 2017). In this study, we mitigated the impact of spatial bias by down-sampling the number of records in certain regions and avoiding rectification for temporal bias because the records were relatively well distributed through the years.

However, several suggestions of data treatment are presented in the literature that are worth considering to further minimise the potential negative impact of the temporal and spatial biases (e.g. Bird et al., 2014; Chapman et al., 2015, Ruiz‐Gutierrez, Hooten, & Grant 2016). Potential improvements in the predictive performance of our approach could also result from the use of ensembles of distinct algorithms over the employment of a single algorithm (BRT, in our case), as observed for the exercises of machine learning classification in other areas (Araújo & New, 2007).

Accompanying the forecasts with measurements of uncertainty adds support to their use for decision-making. In data-driven models the multiple sources of uncertainty and the methods used to measure the magnitude of each have been discussed thoroughly elsewhere (e.g. Buisson, Thuiller, Casajus, Lek & Grenouillet 2010; Ruiz-Gutierrez et al., 2016). Of specific significance to our work is the extent to which the conditions sampled by the occurrences and temporal pseudo-absences represent the environmental combinations being predicted. Given the potentially high-dimensionality of the environmental space, this assessment may not be trivial to evaluate and report. One likely method of overcoming this limitation, as suggested by Kuhn and Johnson (2013), is to first identify and isolate the most important features, and then reduce their dimensionality using techniques such as principal components analysis or multidimensional scaling and finally measure the overlap between the environmental conditions sampled and those predicted in the dimensionally-reduced space. Forecasts made for conditions that either do not overlap with the sample (i.e., extrapolation) or are only sparsely sampled, have higher uncertainty. Besides being useful in the assessment of the reliability of the forecast, the results from this or an equivalent technique, are also relevant in identifying the environmental conditions that will benefit from more intense sampling in future versions of the model.

Our approach would also certainly benefit from being integrated into an ‘iterative ecological forecasting’ framework. Iterative ecological forecasting refers to the continuous updating of the models as new data becomes available (Dietze, 2017; Urban et al., 2016). With the rapidly growing rates at which photographic and non-photographic occurrence records are becoming available, the regular updating of models with the new data may produce a considerable drop in the uncertainty. Notably, such updating would also reduce the uncertainty regarding possible changes in the mechanisms that drive the ecological responses. Changes in the driving mechanisms of the ecological processes can happen even during short time periods (Oliver & Roy, 2015); hence, the use of up-to-date data in the models facilitates lowering the risk of misrepresenting the drivers of the ecological phenomena being forecasted.

The approach we presented in this work does not aim to replace process-based models or correlative models based on large-scale databases of ecological time series. Instead, it aims at being applied to ecological phenomena that cannot be forecasted using these approaches. The employment of occurrence data for spatiotemporal modelling has several conceptual and methodological contingencies, but we have demonstrated that its use for a judicious training of spatial time-series classification models may allow achieving useful forecasts. The approach presented here could be improved in several ways in the future. We expect that investigation on these topics, allied to the continuous increase in the numbers of occurrence records, will help pave the way for a *de facto* contribution towards the forecasting of ecological phenomena.

